# Clearance of senescent cells following cardiac ischemia-reperfusion injury improves recovery

**DOI:** 10.1101/2020.04.28.065789

**Authors:** Emily Dookun, Anna Walaszczyk, Rachael Redgrave, Pawel Palmowski, Simon Tual-Chalot, Averina Suwana, James Chapman, Eduard Jirkovsky, Leticia Donastorg Sosa, Eleanor Gill, Oliver E Yausep, Yohan Santin, Jeanne Mialet-Perez, W Andrew Owens, David Grieve, Ioakim Spyridopoulos, Michael Taggart, Helen M. Arthur, João F. Passos, Gavin D. Richardson

**Affiliations:** Biosciences Institute, Newcastle University, Newcastle upon Tyne, UK; School of Environmental Sciences, Faculty of Science, Agriculture & Engineering, Newcastle University, Newcastle upon Tyne, UK; Faculty of Pharmacy, Charles University, Prague, Czech; School of Medicine, Dentistry and Biomedical Sciences, Centre for Experimental Medicine, Institute for Health Sciences, Queen’s University Belfast, UK; INSERM I2MC, University of Toulouse, France; Translational and Clinical Research, Newcastle University, Newcastle upon Tyne, UK; Department of Physiology and Biomedical Engineering, Mayo Clinic, Rochester, MN, USA

## Abstract

A key component of cardiac ischemia-reperfusion injury (IRI) is the increased generation of reactive oxygen species, leading to enhanced inflammation and tissue dysfunction in patients following intervention for myocardial infarction. In this study we hypothesized that oxidative stress, due to ischemia-reperfusion, induces senescence which contributes to the pathophysiology of cardiac IRI. We demonstrate that IRI induces cellular senescence in both cardiomyocytes and interstitial cell populations and treatment with the senolytic drug navitoclax after ischemia-reperfusion improves left ventricular function, increases myocardial vascularization, and decreases scar size. SWATH-MS based proteomics reveal that biological processes associated with fibrosis and inflammation, that were increased following ischemia-reperfusion, were attenuated upon senescent cell clearance. Furthermore, navitoclax treatment reduced the expression of proinflammatory, profibrotic and anti-angiogenic cytokines, including interferon gamma-induced protein-10, TGF-β3, interleukin-11, interleukin-16 and fractalkine. Our study provides proof-of-concept evidence that cellular senescence contributes to impaired heart function and adverse remodeling following cardiac ischemia-reperfusion. We also establish that post-IRI senescent cells play a considerable role in the inflammatory response. Subsequently, senolytic treatment, at a clinically feasible time point, attenuates multiple components of this response and improves clinically important parameters. Thus, cellular senescence represents a potential novel therapeutic avenue to improve patient outcomes following cardiac ischemia-reperfusion.

## Introduction

Coronary heart disease (CHD) is the leading cause of death and disability in developed countries (1). The most serious manifestation of CHD is ST-segment elevation myocardial infarction (STEMI), which is caused by acute blockage of a coronary artery leading to myocardial ischemia and cardiac cell death. The most effective intervention is timely reperfusion of the myocardium via primary percutaneous coronary intervention (2). Although early intervention can limit acute myocardial infarction (MI) injury, reperfusion can itself induce ischemia reperfusion injury (IRI) resulting in adverse myocardial remodeling and an increased risk of progression to heart failure (3, 4). This necessitates the need to pursue the use of additional therapies to either prevent or ameliorate myocardial IRI, which has been described as a neglected therapeutic target (5).

Cellular senescence is defined as an irreversible cell-cycle arrest characterized by dramatic alterations in gene and protein expression and the production of the senescence-associated secretory phenotype (SASP) (6). The SASP consists of a cocktail of pro-inflammatory cytokines, chemokines, matrix proteases and growth factors that if unhindered can induce many of the biological processes associated with maladaptive cardiac remodeling. These include attenuation of regeneration, induction of fibrosis and cellular hypertrophy and inflammation (7). Furthermore, the SASP can induce senescence in surrounding healthy cells leading to the spreading of senescence throughout affected tissues (8). Senescence can be induced by a variety of stresses including oxidative stress (9, 10).

A key component of IRI is the increased generation of reactive oxygen species (11) which is thought to contribute to tissue dysfunction. Previously, we had shown that cardiomyocyte (CM) senescence can be induced by oxidative stress, accumulates during aging (12, 13) and contributes to age-related myocardial remodeling (12). Furthermore, using the senolytic navitoclax (ABT263), a Bcl-2 family inhibitor, we demonstrated that elimination of age-related senescence prior to myocardial infarction improved outcome in aged mice (13). This data indicates that pre-existing senescence impairs recovery in aged animals, however, the possibility that senescence is induced during IRI and contributes to the disease pathophysiology has not been investigated.

In this study we hypothesized that cellular senescence is an outcome of the oxidative burst occurring during cardiac IRI and is a key contributor to its associated adverse ventricular remodeling and impaired cardiac function. We show that cardiac ischemia reperfusion (IR) induces myocardial senescence and expression of a proinflammatory, profibrotic and anti-angiogenic SASP in young adult mice. Post-IR, treatment with navitoclax clears senescent cells in the heart, attenuates SASP-mediated inflammation, increases myocardial vascularization, reduces scar size and leads to improved cardiac function. Our work provides proof-of-concept that targeting senescent cells may represent a new therapeutic avenue following cardiac IR.

## Results

### Myocardial infarction with reperfusion induces senescence in multiple cardiac lineages

To determine whether senescence is induced in a clinically relevant model of cardiac IRI, 3-4 month-old male mice were subjected to 60 minutes LAD-ligation followed by reperfusion and hearts were collected at different time-points after reperfusion (**Figure 1A**). The qRT-PCR analysis in hearts collected at 4-weeks post-IR demonstrated that mRNAs encoding senescence-associated markers p16 and p21 were increased in the myocardium of IR injured mice compared to controls (**Figure 1B**). Furthermore, histological analysis showed an increased expression of senescent markers both within the infarct and in the peri-infarct region of the left ventricular myocardium (**Figure 1C-E and Supplementary Figure 1A**). At 72 hours, cells within the infarct and the infarct border zone showed increased Senescence-Associated β-galactosidase activity (SA-β-Gal) (**Figure 1C**). SA-β-Gal activity was also evident 1-week post-IR throughout the infarct in interstitial cells and CMs (**Figure 1C and Supplementary Figure 1B**). IRI is associated with increased oxidative stress (14), and in line with this, we saw an increase in 4-HNE staining (a marker of lipid peroxidation) in the infarct zone at 72 hours following IR, with the area demonstrating the highest levels of oxidative stress also displaying the highest senescence burden (**Supplementary Figure 2**). In addition, we found a significant increase in the frequency of cells expressing p16 protein following IR compared with controls (**Figure 1D**). This included CMs, identified via co-expression of troponin-C (trop-C), and cells of the interstitial population. During aging, senescence in CMs is characterized by the presence of telomere-associated DNA damage foci (TAF) which can be induced by oxidative stress (12). To ascertain if telomere dysfunction occurred in CMs following IR, we quantified TAF in CMs within the peri-infarct region. We found that the frequency of CMs positive for ≥5 TAF was significantly increased in IR injured mice (1.53%±0.45 vs 17.71%±7.67 **Figure 1E**).

**Figure 1.**
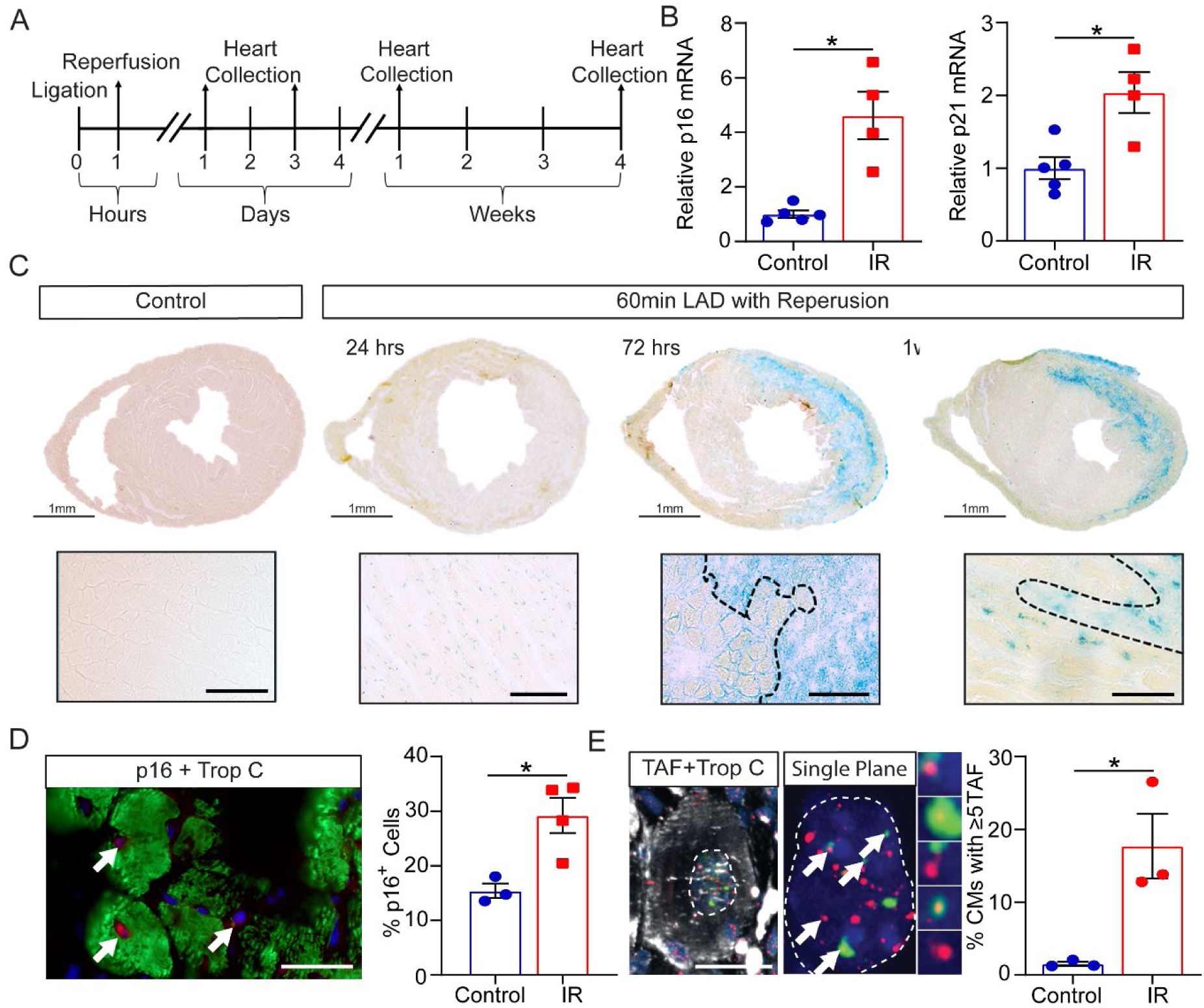
Cardiac ischemia reperfusion induces cellular senescence. **A)** Experimental design. Mice were either subjected to 60 minutes LAD-ligation followed by reperfusion (IR) or received no injury (Control) and hearts collected at the indicated times. **B)** At 4-weeks post-IR hearts were collected and the LV region apical to suture isolated. Real-time qPCR gene expression analysis performed to determine the relative expression of p16 and p21 mRNA (normalized to GADPH). N≥4/group. **C)** SA-β-Gal staining in control and at 24 hours, 72 hours, and 1-week post-IR. **Lower panels)** Higher magnification images of the infarct border. Scale bars 50µm **D)** Quantification of the percentage p16 expressing cells at 4 weeks post-IR. N≥3/group. **Right)** Representative image of p16 expression in trop-C^**+**^ CM (p16 red, trop-C green, DAPI blue. Arrows indicate p16 expressing cells. Scale bar 20µm **E)** Representative images of TAF, γH2AX immuno-FISH, in a trop-C^**+**^ CM (trop-C white, telo-FISH red, γH2AX green). Images are Z-projections of 10μm stacks. Scale bar 10µm. **Middle**) Panel represents a single Z-plane containing TAF (co-localization of a γH2AX foci and a telomere indicated by arrow) and examples of 5 individual TAF **Right)** The mean number of CMs with ≥5-TAF at 4 weeks post-IR. N=3/group. Data are mean±SEM, **B, D**, and **E** were analyzed by 2-tailed unpaired t test; *p<0.05.

### Navitoclax eliminates senescent cells and improves cardiac function following cardiac ischemia reperfusion injury

To ascertain if the observed increase in senescence following cardiac IR contributes to detrimental pathophysiology and represents a valid therapeutic target to improve outcome, we investigated the effects of navitoclax treatment. At 4 days post-IR, once senescence was established, mice were treated with either vehicle only or navitoclax for 7 consecutive days (**Figure 2A**). Immunohistochemical analysis of the LV at 1 month after navitoclax treatment showed a significant reduction in the proportion of p16^**+**^ CMs (24.56%±2.45 vs 6.60%±2.16) and p16^**+**^ interstitial cells (19.76%±1.82 vs 8.90%±2.66) in the peri-infarct region of navitoclax-treated compared to vehicle-treated hearts (**Figure 2B and C).** A reduction in p21 expressing CMs in the peri-infarct region (20.00% ± 1.57 vs 7.12% ± 0.94) was also observed at the same time-point (**Figure 2D and E**). CMs with more than 5 TAF were also reduced in the peri-infarct region in navitoclax treated animals (14.22% ± 2.34 vs 6.41% ± 1.63, **Supplementary Figure 3**). Having demonstrated the effectiveness of navitoclax to reduce senescence, we next aimed to determine if there is a causal relationship between senescence and impaired cardiac function following IR. In a separate cohort of mice, cardiac MRI was performed at 5 weeks post-IR in navitoclax and vehicle-treated mice. All ligated mice demonstrated increased LV end-systolic volumes (ESV) and a decrease in ejection fraction (EF) **(Figure 2F-H)**. Mice treated with navitoclax showed a significantly higher EF than vehicle controls (**Figure 2G**) as a result of a better maintained LV systolic volume (**Figure 2H**). In addition, navitoclax treated mice displayed a significantly larger cardiac output and stroke volume compared to the vehicle group (**Supplementary Figure 4**). No significant difference in EDV was observed between any experimental group (**Figure 2H**). These data support a role for senescence driven myocardial dysfunction following cardiac IR.

**Figure 2.**
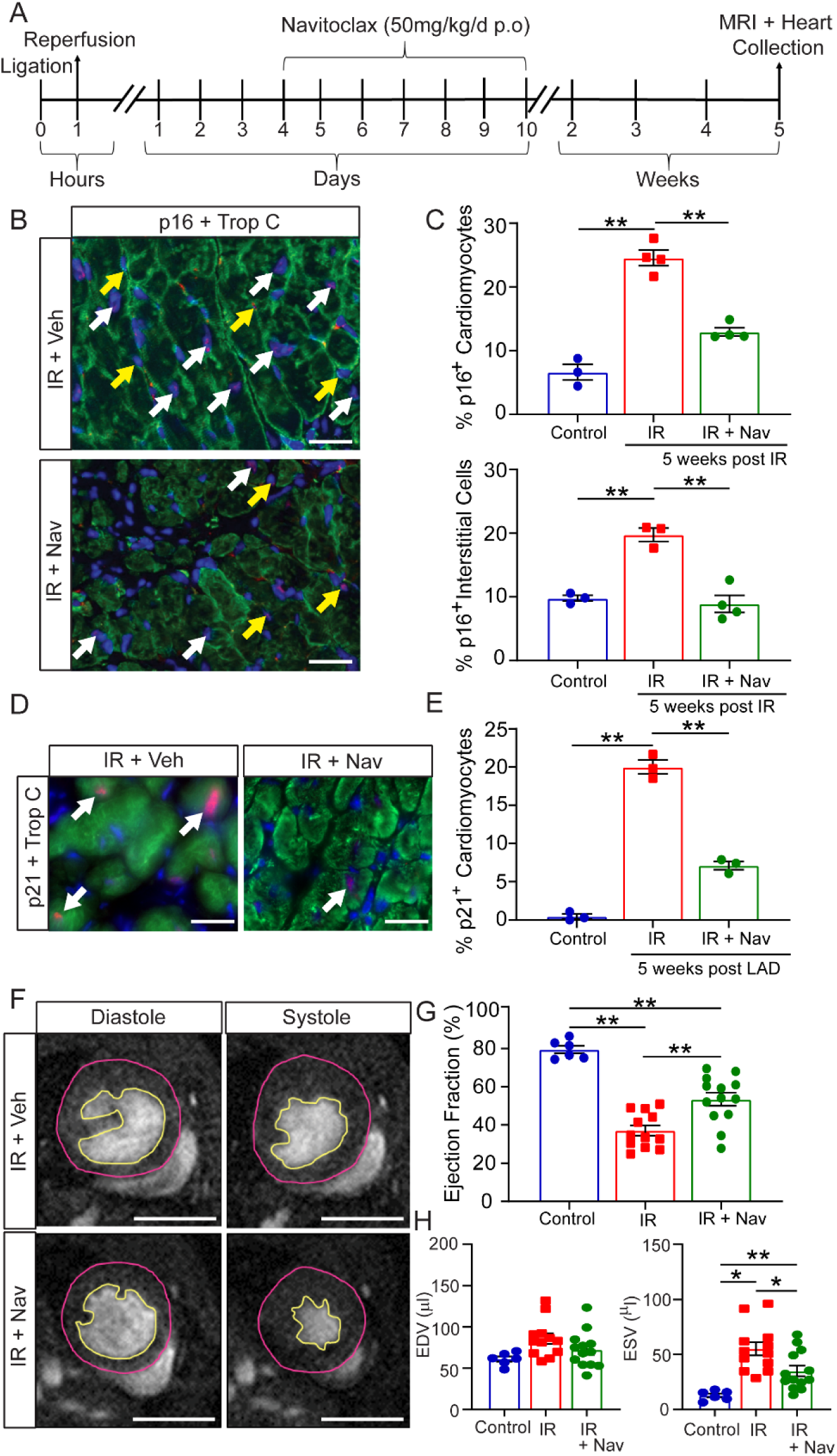
Pharmacological clearance of senescent cells with navitoclax (ABT 263) reduces cellular senescence and improves cardiac ischemia reperfusion. **A)** Mice were either subjected to 60-minute IR or received no injury (Control). After IR mice were randomly assigned to the vehicle (IR) or navitoclax (IR+Nav) groups and provided with navitoclax or vehicle daily from 4 days post-IR for 7 days. MRI was performed at 5 weeks post-IR and hearts were collected. **B)** Representative images of trop-C^**+**^ CMs co-expressing p16 (examples indicated by white arrows) and p16 expressing intestinal cells (examples indicated by yellow arrows) in the LV peri-infarct region (p16 red, trop-C green and DAPI blue). Scale bars 20µm. **C)** Quantification of the percentage trop-C^**+**^ CMs and intestinal cells expressing p16^**+**^ in the LV peri-infarct region. N≥3/group. **D)** Representative images of trop-C^**+**^ CMs co-expressing p21 in the LV peri-infarct region (p21 red, trop-C green, and DAPI blue). White arrows indicate examples of p16^**+**^ CMs. Scale bars 20µm. **E)** Quantification of the percentage of trop-C^**+**^ CMs co-expressing p21^**+**^ in the LV peri-infarct region. N=3/group. **F)** Examples of individual short-axis cine-MR images of mouse hearts. Scale bars 5 mm. **G and H)** EF%, EDV and ESV were calculated based on manual measurements of LV epicardial and endocardial borders. Measurements were made in all cine slices at end-diastole and end-systole. Graphs representing data obtained from MRI analysis. Control N=6, IR N=12 and IR+Nav N=13. Data are mean±SEM. **C, E, G** and **H** were analyzed by one-way ANOVA followed by Tukey’s post hoc test, **p<0.01; *p<0.05.

### Proteome profiling indicates that modulation of molecular pathways involved in remodeling, inflammation and respiration underlies navitoclax mediated improvement in cardiac function post-IR

We next conducted proteomic profiling to identify the protein networks modified by IR and senescence clearance to identify those that may contribute to the observed improvement in cardiac function following treatment. To capture the molecular changes that occur as a result of navitoclax treatment, we performed this analysis at 7 days post-IR (**Figure 3A**). At this time-point, a reduction in p21 expression was observed in the LV of navitoclax treated animals compared to vehicle controls, consistent with elimination of senescence (**Supplementary Figure 5**). Having demonstrated that navitoclax was reducing senescence by day 7 post-IR, LC-MS/MS analysis was performed on protein lysates obtained from LV tissue of vehicle control and navitoclax treated mice at 7 days post-IR and comparable regions of LV from naïve, age-matched control mice **(Figure 3B)**. From a total of 3213 proteins quantified in these LV tissues, 162 were increased (T-test, FDR<0.05) and 142 were reduced following IR. To identify the proteins/pathways modulated by navitoclax selected protein abundance profiles were analyzed to identify proteins increased by IR but attenuated following navitoclax treatment (profile1) **(Figure 3C**). 137 proteins were identified that followed this profile at an FDR <0.01 and 376 at an FDR of <0.05 (**Data set 1**). To assess the potential significance of these proteins to biological functions, pathway analysis was performed in string V 11.0,(15) for the 137 proteins matching the reference with FDR <0.01 **(Supplementary Figure 6)**. Enriched GO terms (biological processes) included processes related to inflammation, such as secretion by cell, cellular secretion, immune response, and response to cytokine (**Figure 3D**). Additionally, the analysis indicated that navitoclax treatment attenuated proteins involved in biological processes related to supramolecular fiber organization and cytoskeleton organization.

**Figure 3.**
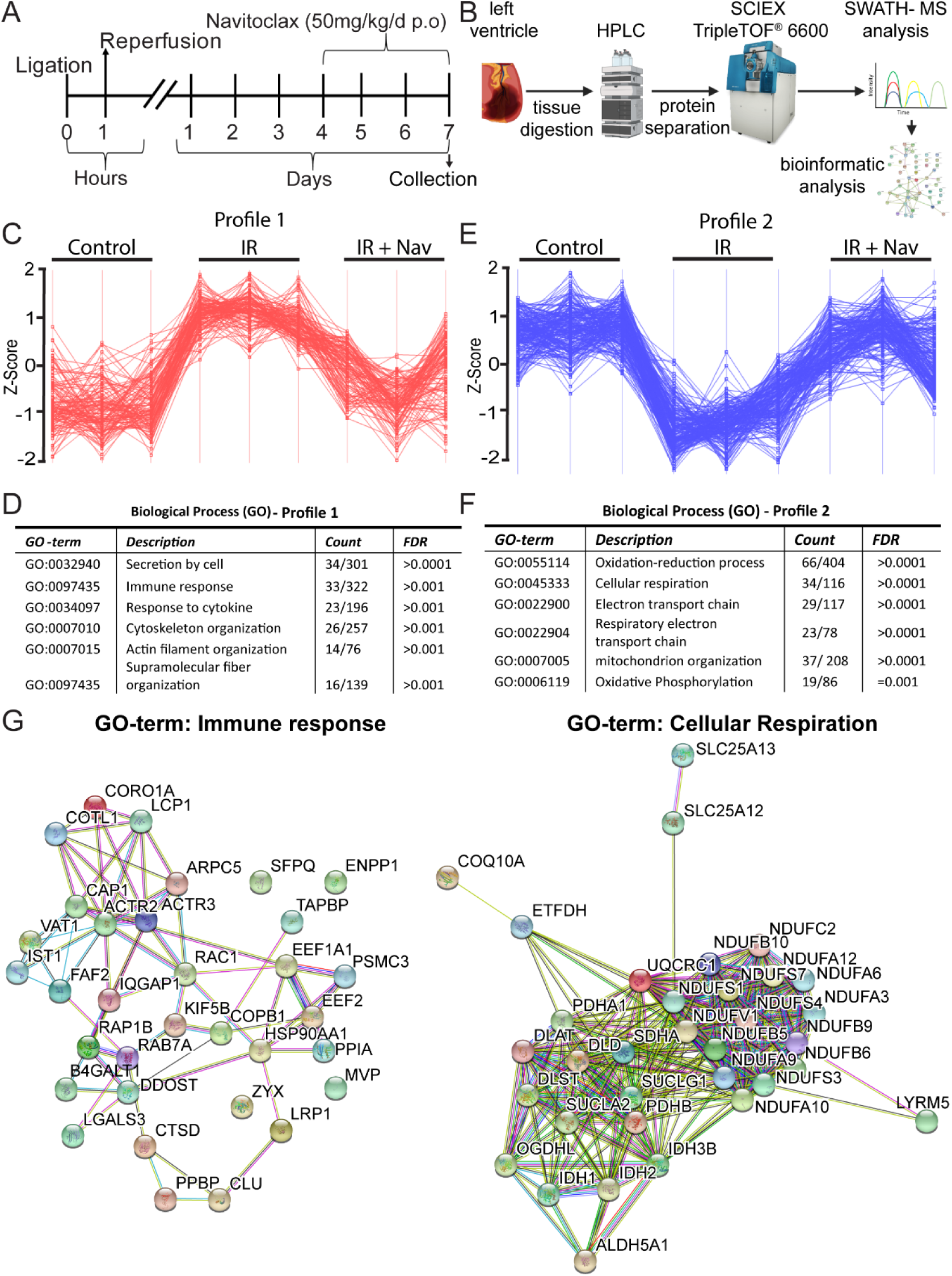
Proteomic pathway profiling. **A)** Experimental design. Mice were either subjected to 60 minute IR or received no injury (Control). After IR mice were randomly assigned to the vehicle (IR) or navitoclax (IR+Nav) groups and provided with navitoclax or vehicle daily from 4 days post-IR for 4 days. On the fourth day of treatment hearts were collected and the LV apical to the suture isolated for analysis (N=3/group). For control mice, a comparable region of LV was isolated for the analysis. (N=3) **B)** Workflow for SWATH-MS proteomics. **C)** Perseus software was used to identify proteins that followed Profile 1. Vertical lines represent biological repeats in each of the indicated groups. Horizontal lines indicate individual proteins that followed the profile at an FDR <0.01. **D)** List of selected significantly enriched GO biological terms for the proteins identified to follow profile 1 at an FDR<0.01. **E)** Perseus software was used to identify proteins that followed Profile 2. Vertical lines represent biological repeats in each of the indicated groups. Horizontal lines indicate individual proteins that followed the profile at an FDR <0.01. **F)** List of selected significantly enriched GO biological terms for the proteins identified to follow profile 2 at an FDR<0.01. **G)** Examples of enriched protein networks for the GO term “immune response”, which followed profile 1 (increased following IR and attenuated by navitoclax treatment), and the GO term “cellular respiration” which followed profile 2 (decreased following IR and rescued by navitoclax treatment).

Next, we aimed to identify the proteins/pathways that followed profile 2 (decreased by IR and rescued following navitoclax treatment) **(Figure 3E**) and identified 199 proteins that followed this profile at an FDR <0.01 and 527 at an FDR of <0.05 (**Data Set 2**). Pathway analysis of the 199 proteins with an FDR <0.01 **(Supplementary Figure 7)**, identified GO terms enriched for this profile were related to cellular respiration and mitochondrial function including, oxidative phosphorylation and the electron transport chain (**Figure 3F)**, suggesting that navitoclax treatment may improve in mitochondrial function and attenuate the oxidative stress caused by IR. Examples of protein networks for the GO term “immune response”, enriched in the proteins following profile 1 and “cellular respiration”, enriched in the proteins following profile 2, are shown in **Figure 3G**.

### Navitoclax attenuates the SASP

Pathway analysis of the proteomics data indicated a reduction of senescence resulted in the modulation of multiple protein networks associated with inflammation and cytokine activity. As such, it is attractive to hypothesize that elimination of senescent cells post-IR improves outcome as a result of attenuation of a pro-inflammatory and profibrotic SASP. To investigate this further a cytokine array was used to evaluate cytokines released within the LV of naïve control and IR hearts treated with either vehicle or navitoclax harvested at 7 days post-injury, as in **(Figure 3A)**. Comparing cytokines present in LV myocardial tissue from vehicle-treated IR and control mice revealed that IR caused an up-regulation of SASP proteins including Interleukin-6 (IL-6), Interferon gamma-induced protein 10 (IP-10), Eotaxin and members of the TGF-β superfamily. Importantly, these SASP proteins were reduced in the navitoclax treated IR animals **(Figure 4 A and B)**. In addition, interleukin-11 (IL-11), interleukin-16 (IL-16), CCL22 and MIP-3β which have been previously associated with cardiac fibrosis, (16, 17) or cardiovascular disease (18, 19), and fractalkine (CX3CL1), a chemokine associated with poorer cardiac functional outcome and increased mortality in MI patients (20), showed a similar trend of reduction following navitoclax treatment. A complete list of cytokines and their expression levels is included in **Supplementary Table I**.

**Figure 4.**
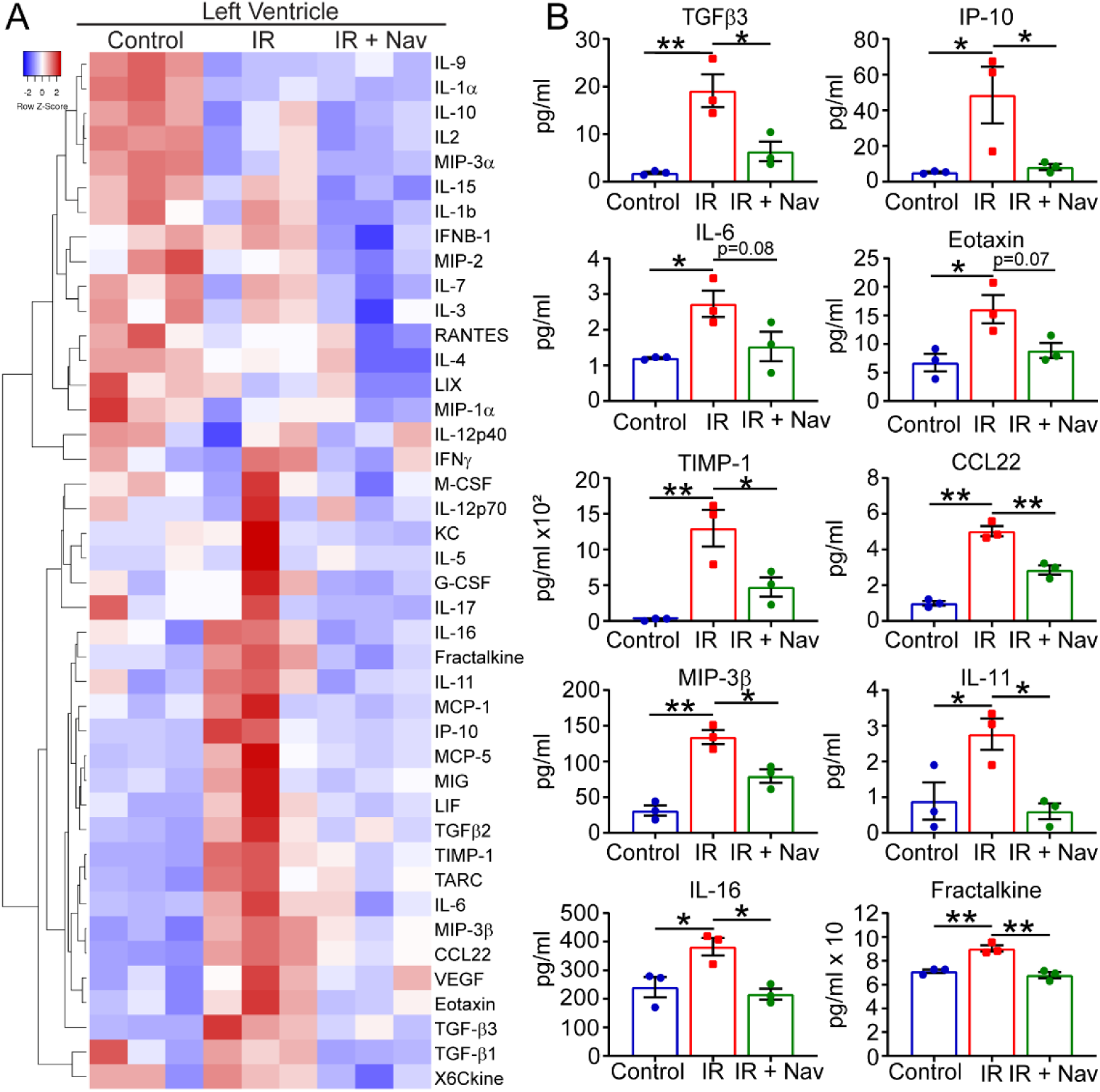
Navitoclax following IR attenuates the SASP. **A)** Clustered heatmap showing all analyzed cytokine protein levels in the LV of naïve control (Control) and IR with either vehicle (IR) or navitoclax treatment (IR + Nav). **B)** Expression of individual protein levels in the LV myocardium of heart in the indicated experimental groups. Data are mean±SEM for each treatment group. N=3/group. Analysis for all proteins was by one-way ANOVA followed by Tukey’s post hoc test, *P<0.05, **P<0.01.

### Navitoclax treatment reduces infarct size and promotes angiogenesis but not cardiomyocyte proliferation in vivo

Navitoclax treatment post-IR reduces expression of SASP proteins with established roles in myocardial remodeling and attenuates biological pathways related to inflammation, ECM production and cytoskeletal organization. These findings led us to hypothesize that elimination of senescence and its associated SASP leads to improved cardiac recovery via a reduction in adverse myocardial remodeling. Mice were treated as previously and also provided EdU to allow quantification of proliferation **(Figure 5A)**. In line with our hypothesis, scar size measured by Masson’s trichrome staining, was significantly reduced in the navitoclax treated mice compared to vehicle control at 5 weeks post-IR (12.47%±1.68 vs 18.50%±2.72 **Figure 5B**). IR also resulted in a significant increase in CM size, however, no difference was observed between the navitoclax treated and vehicle groups **(Figure 5C)**. Accumulation of senescence and expression of the SASP could also impact on regeneration via the bystander effect. To investigate *de novo* CM regeneration following navitoclax treatment we quantified EdU-positive cells in combination with the CM marker trop-C and cell membrane marker wheat germ agglutinin (WGA). We found a trend towards an increase in EdU labelled CMs following navitoclax, however, this was not significant (0.91±0.30 cells per FOV vs 0.72 ±0.14 cells per FOV **Figure 5D**).

**Figure 5.**
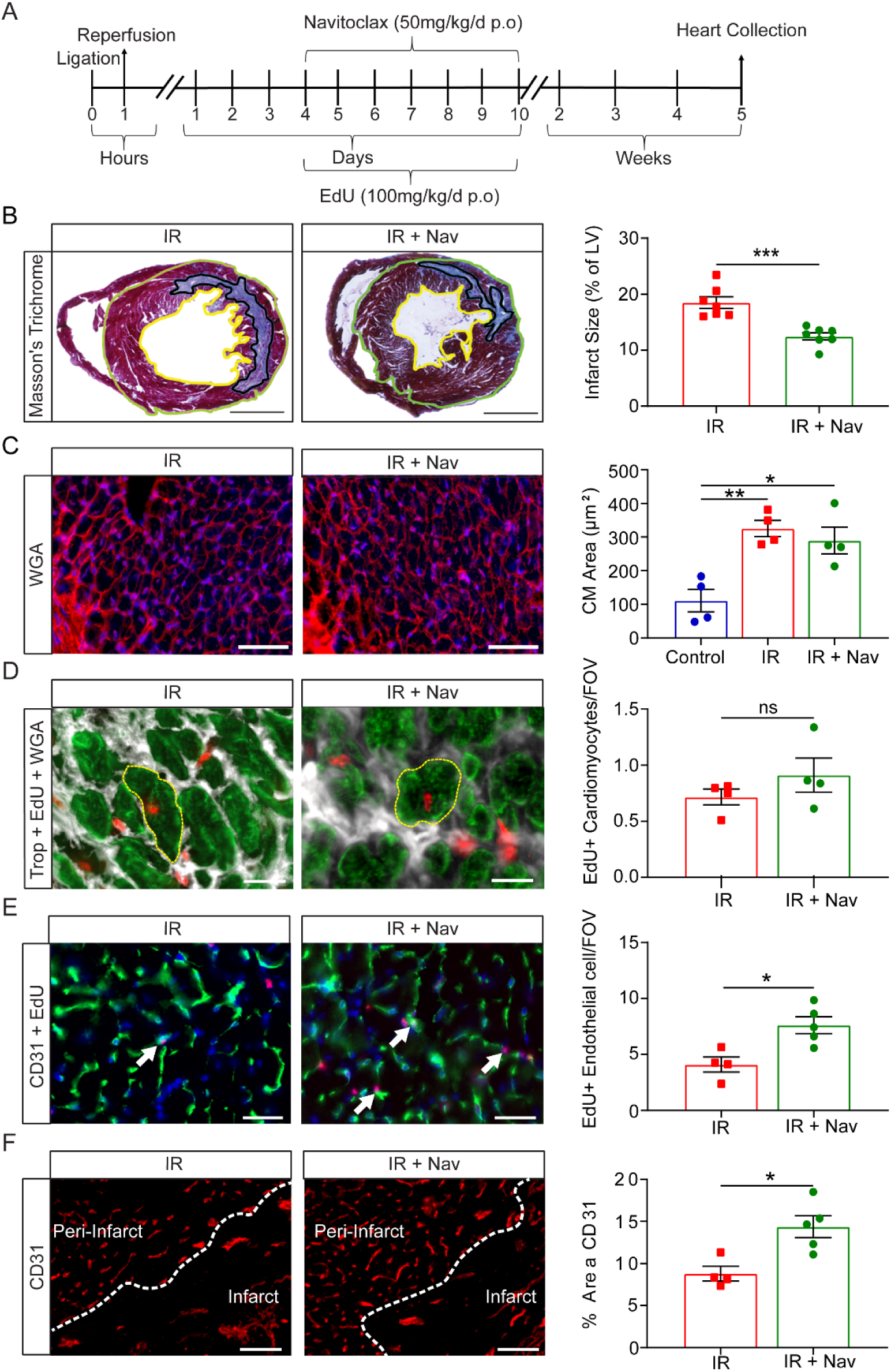
Navitoclax reduces scar size and increases angiogenesis but has no influence on hypertrophy or cardiomyocyte proliferation. **A)** Experimental design. Mice were subjected to 60 minute LAD-Ligation with reperfusion and then at day 4 post-IR were treated with vehicle (IR) or navitoclax (IR + Nav) daily for 7 days. Mice were also provided with EdU during the same period. Hearts were collected at 5 weeks post-IR. **B)** Representative image of Masson’s trichrome staining. Scale bars 1mm. **Right)** Quantification of infarct size relative to total LV area. N=7/group. **C)** Representative images of WGA staining for quantification CM cross-sectional area. Scale bars 50µm **Right)** CM cross-sectional area μm^2^. N=4/group, >150 CMs analyzed per mouse. **D)** Representative image of trop-C^**+**^ CMs co-labelled with EdU highlighted in yellow (Trop-C green, EdU Red and WGA white). Scale bars 20µm **Right)** Quantification of trop-C^**+**^/EdU^**+**^ cells per field of view in the peri-infarct region. N=4/group. **E)** Representative image of CD31^**+**^ expressing endothelial cells co-labelled with EdU (trop-C green, EdU Red). White arrows indicate co-labelled cells. Scale bars 20µm. **Right)** Quantification of the total number of CD31^**+/**^EdU^**+**^ cells per field of view in the peri-infarct region. N≥4/group. **F)** Vessel density in the peri-infarct and infarct zone analyzed using CD31 immunostaining. Scale bars 50µm. **Right)** Quantification of the percentage area of CD31 expression. N≥4/group. Data are mean±SEM, **B, D, E** and **F** were analyzed 2-tailed unpaired t test **C** was analyzed by one-way ANOVA followed by Tukey’s post hoc test, ***p<0.001; **p<0.01; *p<0.05.

Proteomic data analysis also indicated that the elimination of senescent cells and the SASP is associated with improved respiration. This together with the documented anti-angiogenic activities of SASP proteins, including IP-10 (21), led us to further hypothesize that subsequent to IR SASP inhibits endothelial proliferation leading to reduced angiogenesis. Accordingly, a navitoclax mediated reduction in senescence and SASP would attenuate this effect and allow for increased angiogenesis. Histological analysis of EdU together with the endothelial cell marker CD31 showed a significant increase in endothelial cell proliferation in the navitoclax treated animals compared with vehicle controls (7.62±1.67 cells per FOV vs 4.11±1.3 cells per FOV **Figure 5E**). To establish if this could be a result of SASP driving endothelial cells to senescence, we performed *in vitro* conditioned media experiments. Conditioned medium isolated from senescent fibroblasts reduced cardiac endothelial cell proliferation (measured by Ki67 expression and cell output) and increased endothelial superoxide production, a characteristic of endothelial cell dysfunction (**Supplementary Figure 8**). Furthermore, endothelial cells treated with conditioned media from senescent cells demonstrated increased p21 but not p16 expression (**Supplementary Figure 8F**). Finally, we quantified peri-infarct region vascularity. Consistent with an *in vivo* pro-angiogenic effect (22), the peri-infarct region of hearts from animals treated with navitoclax had significantly increased vessel density in the peri-infarct zone compared vehicle control (14.38%±2.88% vs 8.87%±1.74 **Figure 5F**). Taken together our data indicate that in young animals cardiac IR drives senescence and a pro-inflammatory, anti-angiogenic and profibrotic SASP. Moreover, navitoclax mediates the reduction in senescence and improves cardiac function by attenuating SASP reducing scar size and enabling increased angiogenesis.

## Discussion

Efforts to target the oxidative insult that occurs immediately following IR using therapies such as antioxidants have proven unsuccessful (23). The current study provides novel data demonstrating that navitoclax mediates elimination of the senescence that occurs downstream of oxidative stress and that this reduces scar size, increases angiogenesis and improves cardiac function.

Subsequent to cardiac IR, the degree to which remodeling occurs is not only dependent on the immediate detrimental processes associated with ischemia and rapid oxidative stress due to reperfusion but is also dependent on the outcomes of a complex inflammatory response (24). Tissue damage caused by IR induces an acute inflammatory response responsible for the elimination of necrotic debris (25). In mice, this initial response begins to abate at around 5 days, being replaced by a reparative and proliferative phase which functions to suppress the acute response and coordinates tissue remodelling (24). While both phases of inflammation are required for successful wound healing they can also contribute to myocardial dysfunction. The acute inflammatory response can expand tissue damage and a severe or prolonged reparative response is associated with pathological scarring fibrosis (25).

Using a SWATH-MS based proteomic approach we have demonstrated that navitoclax mediated reduction of senescence results in the attenuation of multiple biological processes associated with inflammation and immunity. Furthermore, cytokine array analysis identified that this attenuation included reduced expression of cytokines associated with the nuclear factor-κB (NF-κB) signaling pathway including CCL22, IL-6, IL-11, IP-10, eotaxin and fractalkine (26-31). These proteins are not only typical of the SASP (32) but have also been identified as modulators of the acute inflammatory phase following IR (20). Mouse models in which the NF-κB signaling pathway or individual NF-κB mediated proteins are reduced demonstrate decreased pathological remodeling, a reduction in scar size and improved vascularization (21, 25, 33). Conversely, overexpression of NF-κB associated cytokines, such as IL-6, exhibit adverse remodeling and heightened myocardial inflammation (34). Our data suggest that senescence and the SASP contribute directly to acute inflammation post-IR and senolytics, such as navitoclax, provide a means to attenuate the production of multiple cytokines and chemokines, known to be detrimental to recovery.

As senescence was evident by 3 days post-IR, we treated mice from this time point using a dose of navitoclax that we have previously shown to induce myocardial senolysis (12). This treatment extended into the reparative phase of the response to heart injury, a phase that is characterized by the secretion of TGF-β1, an anti-inflammatory and pro-fibrotic cytokine (35). TGF-β expression is dynamic in the heart post-IR, where it is critical for the switch from the pro-inflammatory to the resolution phase of cardiac healing, and for driving formation of the fibrotic scar. In mice, TGF-β1 and β2 expression peaks at 6–72 hours post-reperfusion and declines after 3 to 7 days (36, 37). TGF-β3 expression is induced at a later 3-7 day time-point and is maintained at high levels over a longer time frame. Early neutralization of TGF-β signaling at 24h post-MI is detrimental as it increases both cardiac dysfunction and mortality, whereas late disruption of TGF-β signaling is protective for fibrosis and hypertrophic remodelling (38).

Here we demonstrate, as with the aging heart, there is an association between the level of myocardial TGF-β ligand expression and the level of myocardial senescence. It is, therefore, possible that the SASP also contributes to driving fibrosis and scar formation; a hypothesis supported by the reduced scar size observed in the navitoclax treated animals. While our observations initially appear at odds with studies demonstrating that senescence is required to attenuate cardiac fibrosis (39, 40), it is important to note that navitoclax does not inhibit senescence, but induces apoptosis once senescence is achieved (41). Therefore, the two observations are entirely compatible. Indeed Zhu *et al* propose that while fibroblast senescence reduces collagen deposition in the short term post-MI, senescent fibroblasts are also a source of chronic inflammation contributing to ongoing cardiac fibrosis in the longer term (40).

In the absence of MI, aged animals demonstrated an increase in CM renewal following either pharmacogenetic or pharmacological clearance of senescence (12, 42). However, senescence elimination following IR did not enhance CM regeneration in the young animals used in our study. Following MI, both the vehicle and navitoclax treated mice displayed a similar CM regenerative response, corresponding to a renewal of 0.47% and 0.38% CMs, respectively, during the first week following MI. These rates are consistent with those previously reported for MI hearts without therapeutic intervention (43), suggesting that young animals have a CM regenerative potential that is not impeded by senescence. In contrast, we noted an increase in both endothelial proliferation and vessel density in the hearts of navitoclax treated mice. Together with our *in vitro* data showing the anti-angiogenic effects of senescent fibroblasts, and the known anti-angiogenic properties of the SASP (7), the increased angiogenesis post-IR following navitoclax treatment is likely to have occurred via the reduction in SASP. Improved angiogenesis and improved myocardial oxygenation may also contribute to the observed improvement in cellular respiration. In this context, however, we have previously demonstrated that myocardial ageing and accumulated senescence is associated with mitochondrial dysfunction (12). Therefore, a reduction in senescence cells with dysfunctional mitochondria, by navitoclax treatment, may also contribute directly to a global increase in mitochondrial function.

In conclusion, this data suggests a key role of myocardial senescence signaling downstream of the initial oxidative stress response to reperfusion. We suggest that targeting senescence is a valid and clinically feasible strategy to attenuate maladaptive remodeling and promote recovery post-IR. Major advantages of senolytic treatment over current strategies include: 1) targeting senescence which occurs as a result of the cellular stress associated with IRI provides an extended therapeutic window; and 2) targeting senescence attenuates multiple components of the inflammatory responses subsequent to IR, which are detrimental to recovery (24, 25) Recent studies have begun to trial senolytics in patients suffering from pulmonary fibrosis or kidney disease (44, 45), and if senolytics prove to be effective and safe they could be transformative for cardiovascular medicine.

## Methods

### Mouse model and myocardial infarction with reperfusion

Male C57BL/6J mice at 3-4 months of age were used in all studies. Intra-operative analgesia was induced by pre-treating mice with fentanyl/fluanisone (0.4ml/kg, Hypnorm), prior to anesthesia using isoflurane, which was maintained using mechanical ventilation following endotracheal intubation (3% isoflurane/97% oxygen, 130-140 stroke rate, stroke volume initially 5ml/kg – increased to 7.5ml/kg post-thoracotomy). At the fourth-intercostal space, left-side thoracotomy was executed to allow partial removal the pericardium and enable a 7-0 prolene suture to be placed around the left anterior descending artery (LAD) and loosely tied. An infarction was induced by inserting 2mm PE-10 tubing into the suture loop and tightening the suture knot to terminate blood flow for 60 minutes. The tubing was removed to allow myocardial reperfusion, the chest cavity closed and 0.05mg/kg, Vetergesic was provided as analgesia. Naïve mice (no surgical intervention) were used as control.

### In vivo navitoclax (ABT263) treatment

Navitoclax was prepared in a lipid vehicle solution consisting of EtOH, polyethylene glycol 400 and Phosal 50 PG in a 1:3:6 ratio, respectively (46). Mice were randomly assigned to experimental groups. Navitoclax (50mg/kg/day via oral gavage) was provided for the dose timing regimes detailed in the main text. When required, EdU (100mg/kg/d) was provided via intraperitoneal injection (47, 48).

### Histology and Immunohistochemistry

Immunohistochemistry was performed as described previously (49). 10 μm sections were used for all studies. Primary antibodies used: rat ant-p21 (HUGO291, Abcam ab107099), goat-anti-troponin C (Abcam, ab30807), rabbit anti-p16 (Rockland, 100-401-170) and rat anti-CD31 (MEC13.3, BD Biosciences, 550274). Secondary antibodies used were donkey anti-rat AF594 (Life Technologies, A21209), donkey anti-goat AF 488nm (Life Technologies, A11055), donkey anti-rabbit AF594 (Life Technologies, R37119), donkey anti-goat AF488 (Life Technologies, A11055) and donkey anti-mouse AF647 (A31571, Life Technologies). Slides were mounted in Vectarshield containing DAPI (Sigma, MBD0015). 5-ethynyl-2-deoxyuridine (EdU) labelling performed with Invitrogen Click-iT EdU Alexa Fluor 594 Imaging Kit (Life Technologies, C10339).

### Senescence-associated β-galactosidase staining

Cryo-embedded sections were stained using the senescence associated β-galactosidase (SA-β-Gal) staining kit (Cell Signaling Technology, 9860) as per the manufacturer’s instructions with the following modifications for tissue sections. Slides were thawed at room temperature, fixed using the provided fixative solution for 15 minutes and washed three times in PBS. The β-galactosidase staining solution was prepared at pH 6 and added to each slide. Slides were incubated at 37°C for 24 hours. Once developed slides were washed with PBS, dehydrated in 95% and 100% ethanol solutions, washed in Histoclear and mounted with Histomount

### Real-Time PCR (qRT-PCR)

Qiagen RNA extraction kit was used for RNA isolation. First strand cDNA synthesis kit (Thermo Fisher Scientific, Waltham, MA, US) was used for cDNA synthesis. Real-time PCR was performed in a 7500 Fast Real-Time PCR System using Taman probes (Life Technologies Ltd, UK) GAPDH (Mm99999915_g1), p21 (Rasa3) (Mm00436272_m1) and p16 (Cdkn2a) (Mm00494449_m1). Data presented normalized to GAPDH.

### Immuno-FISH

Telomere-associated DNA damage foci (TAF) were detected by performing Immuno-FISH, as previously described (50), on cryo-embedded heart sections. Briefly, sections that were labelled with rabbit monoclonal anti-γH2Ax (20E3, Cell Signaling Technology, 9718) and following secondary labelling with goat anti-rabbit IgG biotinylated (VectorLab, PK-6101), sections were fixed with methanol: acetic acid (3:1), dehydrated and incubated with PNA hybridization mix with 5% blocking reagent (Roche) containing 2.5 μg/ml Cy3-labelled telomere-specific (CCCTAA) peptide nucleic acid probe (Panagene).

### Microscopy and image analysis

All images were acquired using Axio Imager (Zeiss) and analyzed using ZEN 2.3 (Zeiss). The peri-infarct region; was defined as the region proximal to the infarct **(Supplementary Figure 1A, red region)**. CM hypertrophy was quantified as described previously (51), following staining with the membrane marker wheat germ agglutinin (WGA) (W32466, Invitrogen, UK).

### Magnetic resonance imaging and analysis

Magnetic resonance images were acquired using a horizontal bore 7.0T Varian microimaging system (Varian Inc., Palo Alto, CA, USA) equipped with a 12-cm microimaging gradient insert (40 gauss/cm) and analyzed as described previously (22).

### Liquid Chromatography Mass Spectrometry (LC-MS/MS) with SWATH acquisition (Sequential Windowed Acquisition of All Theoretical Fragment Ion Mass Spectra)

#### Left ventricular sample preparation

At the time points detailed in the results section, hearts were collected, the left ventricle (LV) was dissected apical to the suture and placed in RIPA buffer (R0278, Sigma) containing protease inhibitors. Tissue was homogenized and proteins precipitated with acetone. Protein was dissolved in 8M urea, 10mM HEPES at pH 8.0. 200µg of protein was reduced with 30mM DTT at 30°C for 30 minutes followed by alkylation with 10mM Iodoacetamide. Urea molarity was reduced to 1.5M prior to trypsin digestion (Worthington, TPCK treated) at a ratio of 20:1 (protein: trypsin). The digestion was stopped with trifluroacetic acid (TFA) and proteins purified with a self-packed C18 stage.(52) Peptides were eluted, dissolved in 3% acetonitrile with 1% TFA and sonicated.

#### Nano LC-MS/MS

Analysis was performed with an AB-Sciex 6600-Tripletof operating in SWATH mode. Protein digests were injected into a mass spectrometer through an UltiMate 3000 RSLCnano system. Samples were loaded onto a 300μm x 5mm C18 PepMap C18 trap cartridge in 0.1% formic acid and then separated at 300nl/min using a 95min nonlinear gradient (3-30%ACN:87min; 30-40%:10min; 40-90%:5min) using a 75μm x 25cm C18 column (ReproSil-Pur Basic-C18-HD, 3 µm, Dr. Maisch GmbH). The eluent was directed to an Ab-Sciex TripleTOF 6600 mass spectrometer through the AB-Sciex Nano-Spray 3 source, fitted with a New Objective FS360-20-10 emitter. SWATH acquisition was performed in a mass range of 400-800m/z, with 75 variable SWATH bins and accumulation time of 40 milliseconds. The number of SWATH bins was estimated based on a pilot DDA run of a random sample using SWATH variable window calculator ver.1.1 (AB Sciex).

All the raw data with the associated search results were deposited in a publicly accessible repository (https://massive.ucsd.edu/, MSV000085040).

#### Data analysis

The SWATH files were processed in PeakView 2.1 (AB Sciex) with the SWATH MSMSall micro app (top 1000 peptides/protein, top 5 transitions/peptide, confidence threshold: 95%, FDR threshold: 1%, modified peptides excluded, extraction window: 10min, XIC width: 10ppm). The mouse heart spectral library was obtained from ProteomeXchange (53) (PXD017795, MSV000085036) and pre-aligned to a randomly selected SWATH file, as described (54). The results were exported, via MarkerView 1.2.1 (AB Sciex), as tab-separated ASCII files and processed using the Perseus software framework (55). Protein peak areas were transformed to log scale, followed by median subtraction (sample median was subtracted to account for loading differences) and the values derived from technical replicates were averaged. Individual proteins had to be quantifiable in all samples to meet the criteria for further analysis. T-test was applied to identify proteins that differed in relative abundance between experimental groups and permutation-based false discovery rate was estimated to account for multiple comparisons. Protein groups, following the relative abundance profiles of interest, were then extracted. The log2 peak areas were first z-scored for each protein and the reference profile was defined. For each protein, a sum of squared differences (differences between measured values and the reference) was calculated as a measure of distance from the reference. The resulting values were then used for permutation-based FDR calculation **(R-Script file supplementary data)** to allow confident identification of proteins that followed a predefined abundance profile. Association with particular subcellular compartments, biological functions or pathways, was assessed for molecules differentially abundant between conditions and/or sharing the same abundance profile, using STRING v11.0 (15), followed by manual interrogation. Statistical background for pathway/enrichment analysis was restricted only to proteins detected in the analyzed samples.

### Scar size quantification

Masson’s trichrome staining was performed in order to visualize scar tissue (56). Hearts were sectioned into 5 sets of slides, 10 slides per set and stained with Masson’s trichrome. Each set of slides were imaged and analyzed using the Leica Digital Image Hub. The LV area was calculated by measuring the epicardial area and subtracting the endocardial area. The infarct area was then measured and the percentage of LV that is infarct calculated to analyze scar size.

### Cell culture

Cell culture work has been performed using human embryonic lung MRC5 fibroblasts acquired from European Collection of Authenticated Cell Cultures (ECACC), Salisbury, UK and mouse cardiac endothelial cells from Cedarlane (Ontario, Canada).

### Statistical analysis

All analysis was performed in a blinded manner. All analysis was performed using GraphPad Prism 8.0. Data were first tested for normality. Parametric data were analyzed using a 2-tailed t test or one-way ANOVA, as appropriate.

### Study approval

All animal studies were conducted in accordance with the Guidance on the Operation of the Animals (Scientific Procedures) Act, 1986 (UK Home Office), and approved by the local ethics committee.

## Supporting information

Data set 1

Data set 2

r script file

## Author Contributions

ED, AW, RR, PP, ST-C, AS, JC, EJ, LDS, EG, OEY, YS performed experiments. GDR, JFP, JMP, HMA, IS, WAO, MT and DG contributed to supervision. PP, MT and JFP designed and supervised aspects of the study. GDR designed, supervised, and oversaw the study and wrote the manuscript with input from all authors.

## Acknowledgements

This study was funded by; The British Heart Foundation PG/19/15/34269, PG/14/86/31177, PG/18/25/33587 PG/14/86/31177, PG/18/57/33941, Wellcome Trust and the Newcastle Healthcare Charity. JFP would like to acknowledge the Ted Nash Long Life Foundation. Figure 3B was created with BioRender.com.

The authors have declared that no conflict of interest exists

