## Supplementary figures and Tables for "Clearance of senescent cells following cardiac ischemia-reperfusion injury improves recovery"

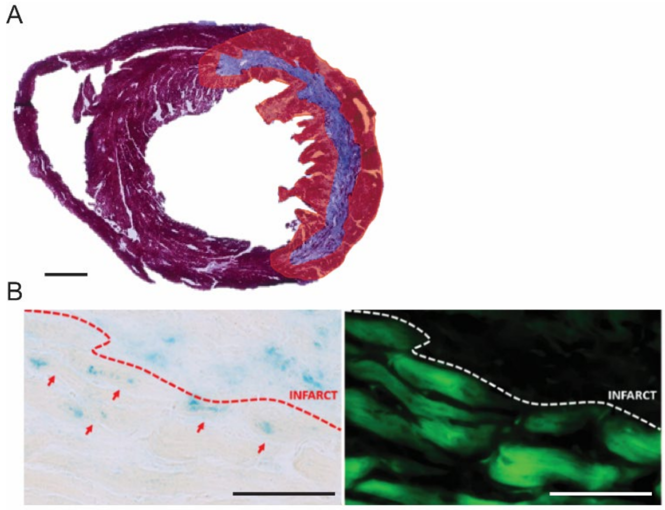


**Supplementary Figure 1. A)** Defined peri-infarct region used for quantification studies. Scale bar 500µm **B)** Representative image of CMs staining positive for SA-β-Gal in the peri-infarct region following IR (blue – SA-β-Gal; green – Auto fluorescence). Scale bar 50µm

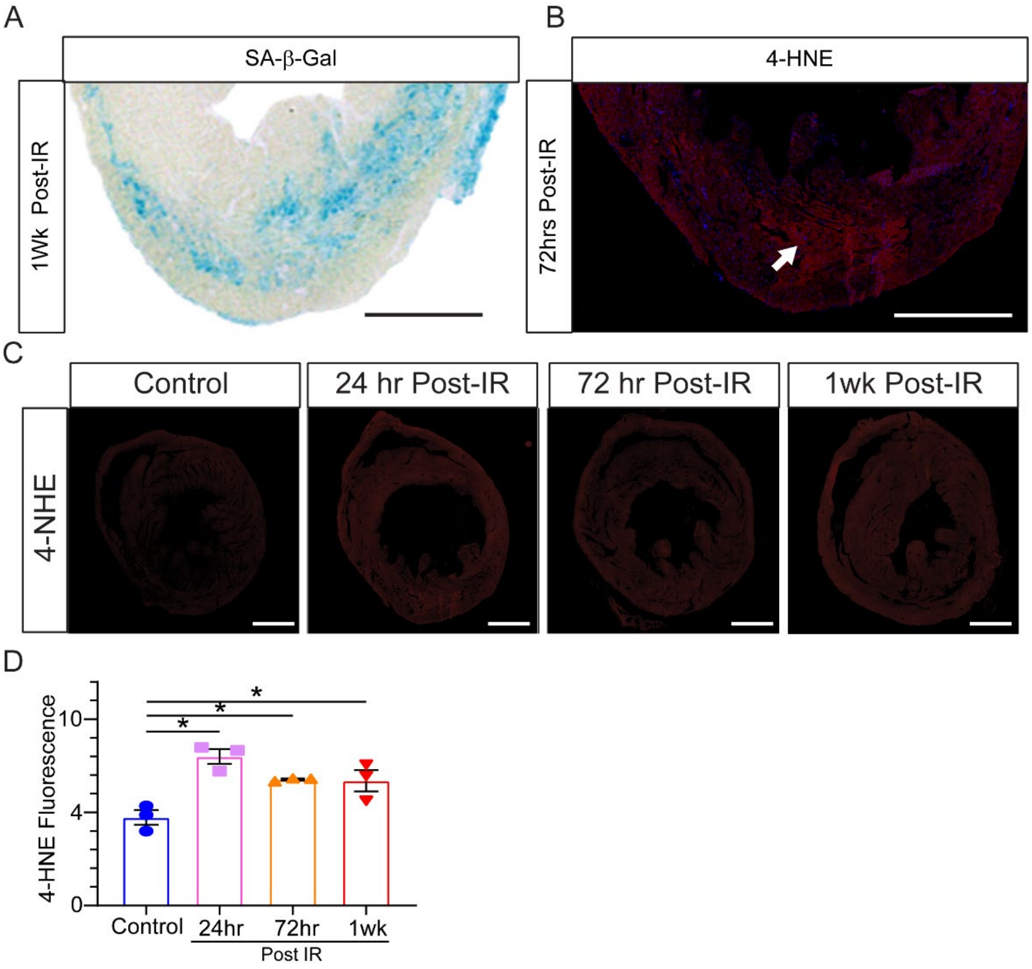


**Supplementary Figure 2. A and B)** Representative image of 4-HNE and SA-β-Gal staining at 72-hours and 1-week, respectively. White arrow indicates area of high 4-HNE in the area of infarct. **C)** Representative image of 4-HNE at each investigated time point post-IR. **D)** Quantification of 4-HNE demonstrates increased myocardial superoxide production in the heart subjected to 60 minute LAD-ligation with reperfusion. N=3/group. All scale bars 2mm. Data are mean±SEM, analysis by one-way ANOVA followed by Tukey's post hoc test, * $P<0.05$.

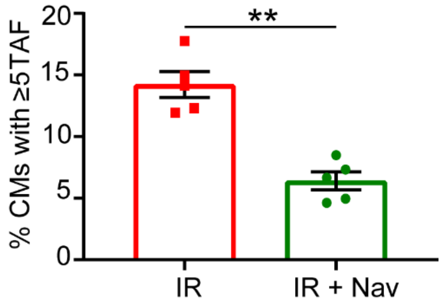


**Supplementary Figure 3.** Quantification of the percentage of ≥5 TAF$^{+}$ CM in the peri-infarct region of the left ventricle peri infarct region in vehicle or Navitoclax treated mice at 5 weeks post-IR. N=5/group. Data are mean±SEM, analysis by 2-tailed unpaired t test, ** P<0.01.

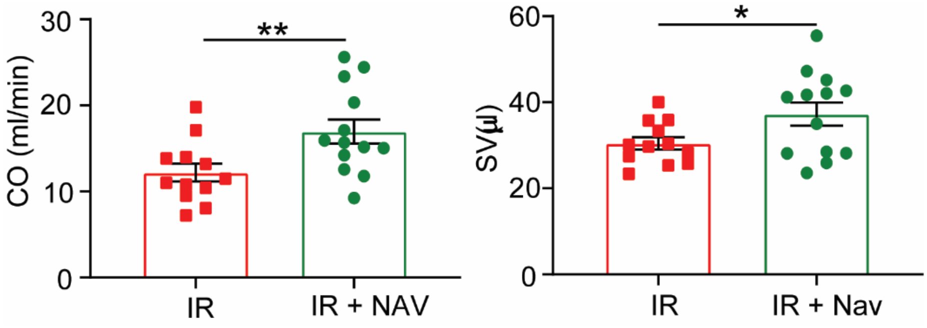


**Supplementary Figure 4.** Data obtained from MRI analysis for mice treated with either vehicle or navitoclax post-IR. IR N=12 and IR+Nav N=13. Cardiac output (CO ml/min) and stroke volume (SV μl). Data are mean±SEM, analysis by 2-tailed unpaired t test **P<0.01.

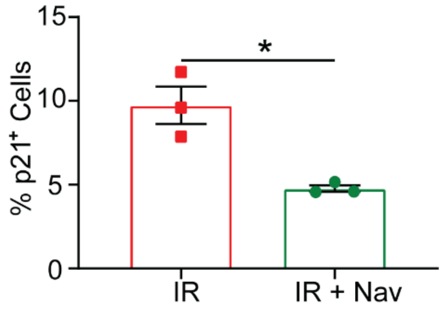


**Supplementary Figure 5.** Quantification of the percentage of p21 expressing cells in the LV peri infarct region of vehicle control and navitoclax treated animals at 7 days post-IR. N>3/group. Data are mean±SEM, analysis by 2-tailed unpaired t test *P<0.05.

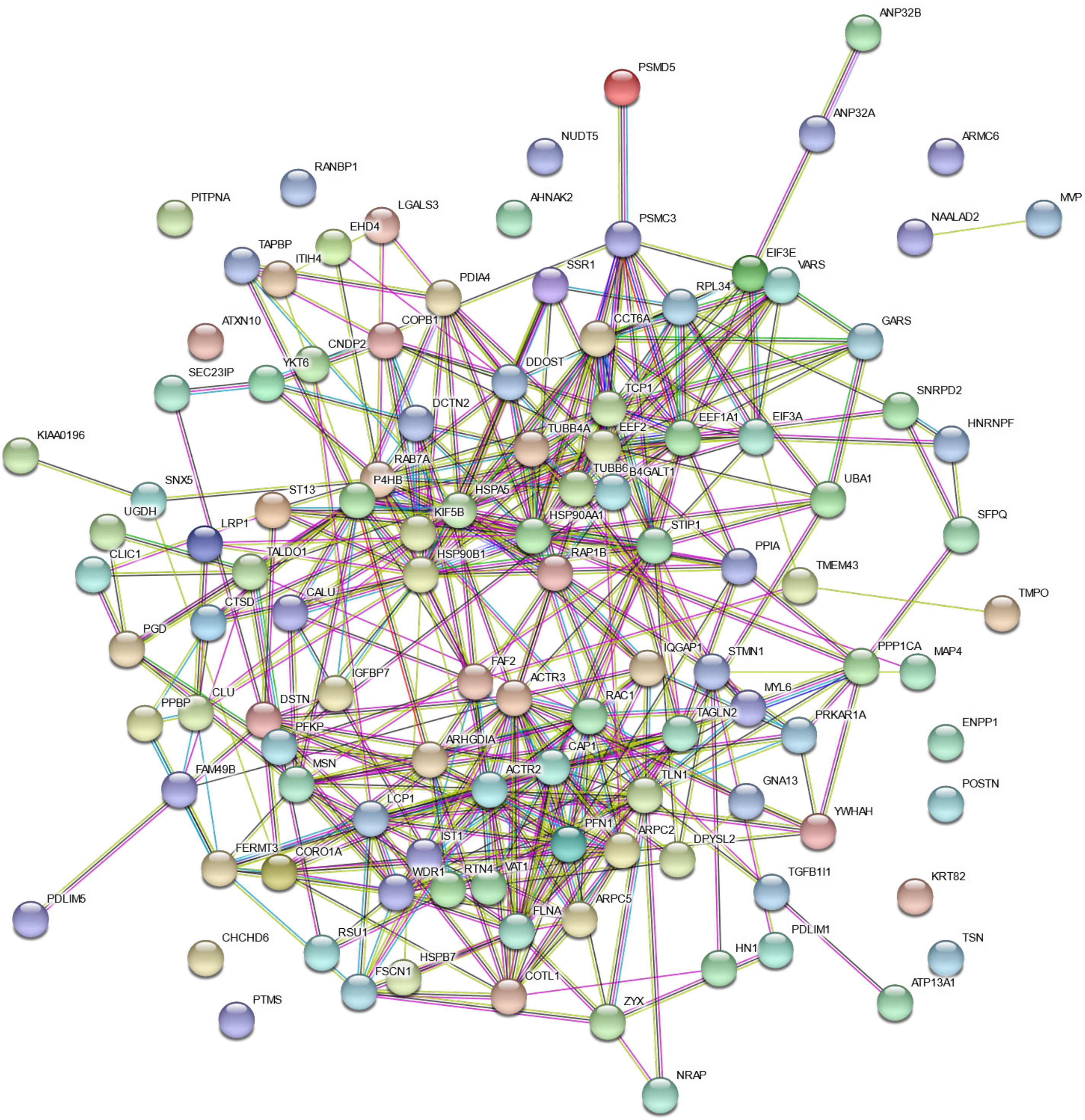


**Supplementary Figure 6.** Protein network for proteins identified for expression profile 1 (increased following IR and reduced after navitoclax treatment).

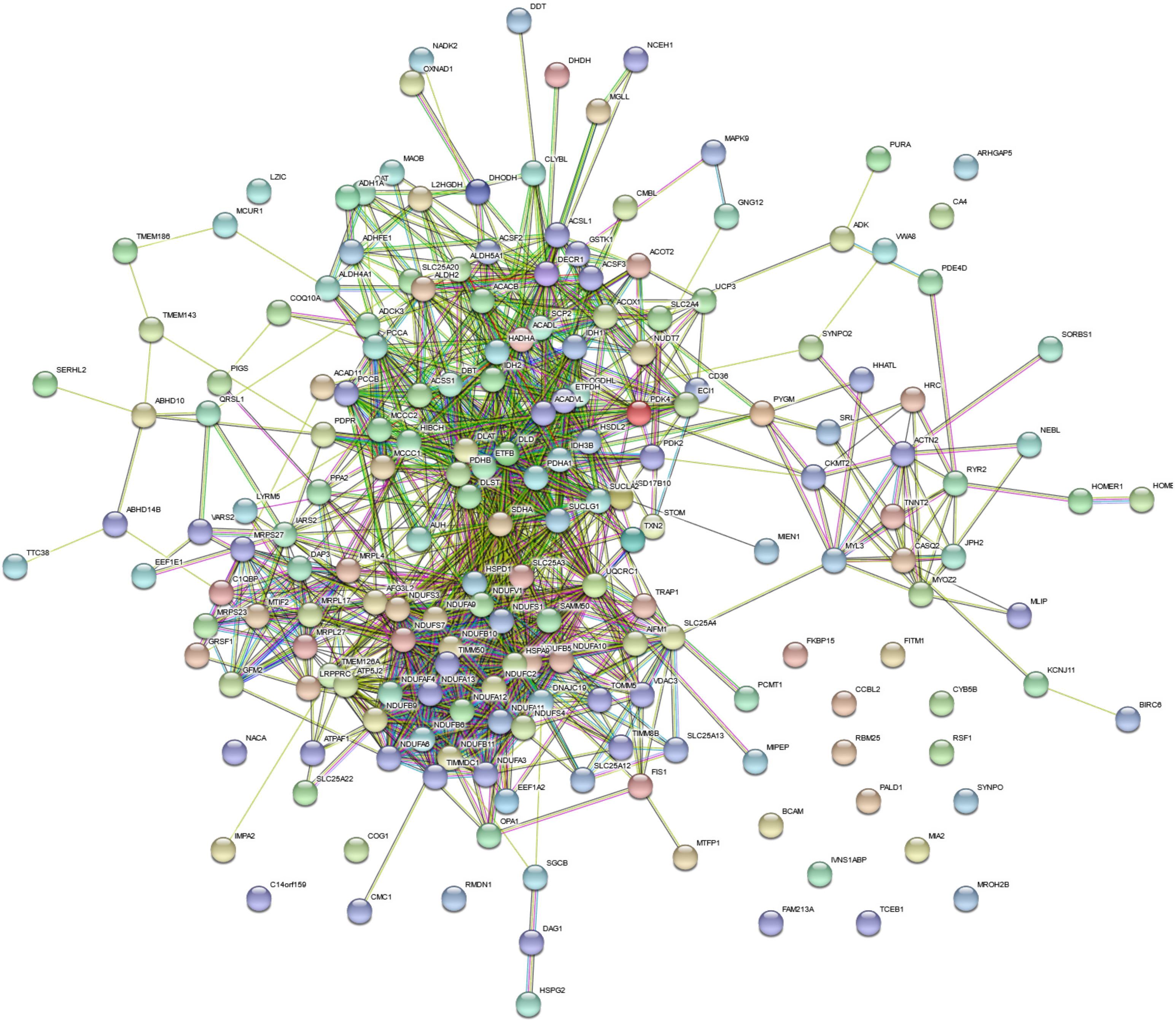


**Supplementary Figure 7.** Protein network for proteins identified for expression profile 2 (decreased following IR and rescued by navitoclax treatment).

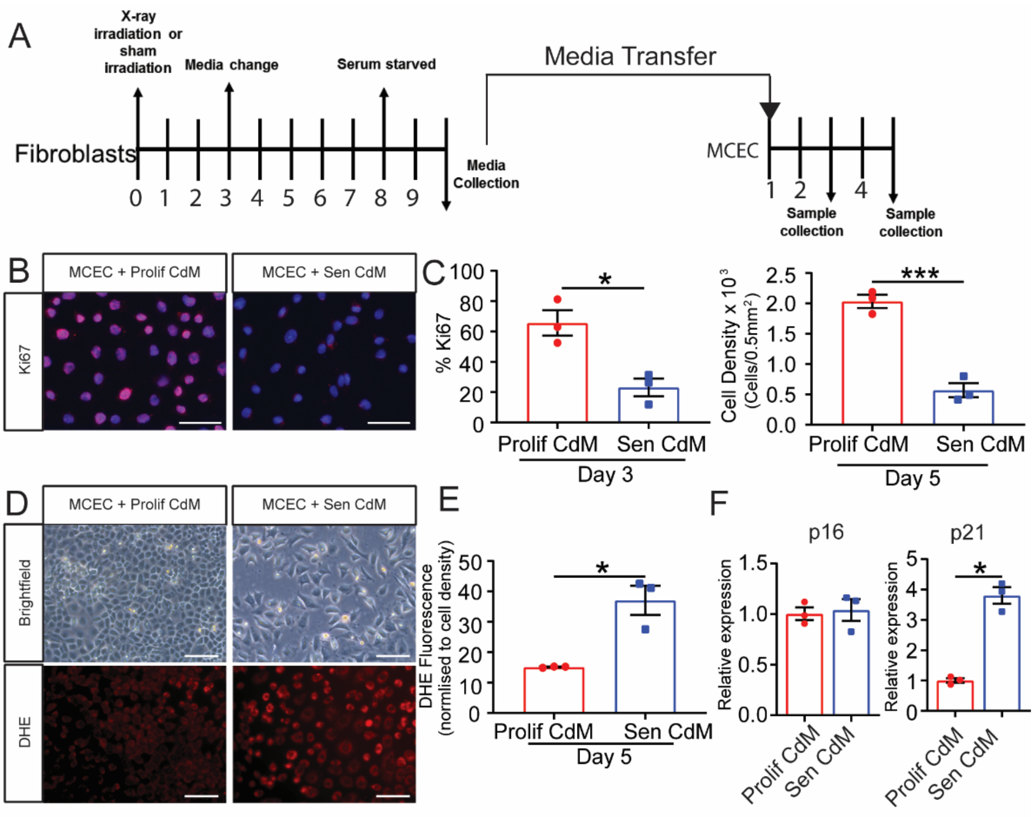


**Supplementary Figure 8. SASP induces reduces proliferation in primary cardiac endothelial cells. A)** Experimental Design, MRC5 fibroblasts were either induced to senescence with X-ray irradiation or sham irradiated. Following 10 days, conditioned medium (CdM) was collected and used to culture mouse cardiac endothelial cells (MCEC). Treated endothelial cells were collected at day 3 and day 5. **B)** Representative images of Ki67 expression in MCEC cultures on day 3. **C)** Quantification of the percentage of MCEC expressing Ki67 at 3 days and MCEC density at 5 days following culture in CdM from either proliferative (Prolif CdM) or senescent fibroblasts (Sen CdM). **D)** Representative bright-field images and images of DHE staining of MCECs at day 5 for each CdM. **E)** Quantification of the in intensity DHE fluorescence normalized to total cell number at 5 days of CdM treatment. **F)** Real-time PCR gene expression analysis for relative p16 and p21 expression (normalized to GAPDH). All scale bars 50μm Analysis by 2-tailed unpaired t test, *P<0.05, **P<0.01, ***P<0.001.

**Supplementary Table 1. Values from MD-44 and TGFB1-3 Arrays.** Data from n=3. Statistical test by one-way ANOVA.

| Cytokine | Concentration (pg/ml ± SD) | | | p Value | | |
|---|---|---|---|---|---|---|
| | Control | IR + Veh | IR + Nav | Control vs IR + Veh | Control vs IR + Nav | IR + Veh vs IR + Nav |
| Eotaxin | 6.74 ± 2.67 | 16.10 ± 4.28 | 8.87 ± 2.29 | 0.0270 (*) | 0.7081 | 0.0728 |
| EPO | - | - | - | - | - | - |
| Fractalkine | 71.21 ± 2.53 | 90.27 ± 4.56 | 67.98 ± 4.49 | 0.0022 (**) | 0.3579 | 0.0014 (**) |
| G-CSF | 0.78 ± 0.17 | 1.17 ± 0.38 | 0.56 ± 0.07 | 0.1981 | 0.5537 | 0.0499 (*) |
| GM-CSF | - | - | - | - | - | - |
| IFNB-1 | 66.52 ± 1.94 | 67.22 ± 1.05 | 60.25 ± 2.88 | 0.9130 | 0.0243 (*) | 0.0153 (*) |
| IFNγ | 0.95 ± 0.79 | 2.09 ± 0.13 | 0.62 ± 0.72 | 0.2532 | 0.8323 | 0.1373 |
| IL-1α | 51.00 ± 6.92 | 17.17 ± 5.02 | 14.14 ± 2.21 | 0.0005 (***) | 0.0003 (***) | 0.7571 |
| IL-1β | 7.20 ± 0.77 | 6.65 ± 1.10 | 5.28 ± 0.36 | 0.6952 | 0.0560 | 0.1619 |
| IL-2 | 25.73 ± 0.52 | 19.78 ± 2.93 | 17.75 ± 1.41 | 0.0200 (*) | 0.0051 (**) | 0.4403 |
| IL-3 | 0.94 ± 0.12 | 0.84 ± 0.10 | 0.67 ± 0.16 | 0.6607 | 0.0969 | 0.3003 |
| IL-4 | 0.19 ± 0.01 | 0.15 ± 0.01 | 0.12 ± 0.05 | 0.2437 | 0.1013 | >0.9999 |
| IL-5 | 0.01 ± 0.00 | 0.05 ± 0.00 | 0.01 ± 0.00 | - | - | - |
| IL-6 | 1.21 ± 0.04 | 2.73 ± 0.64 | 1.53 ± 0.71 | 0.0347 (*) | 0.7631 | 0.0841 |
| IL-7 | 3.57 ± 0.46 | 2.93 ± 1.02 | 1.74 ± 0.42 | 0.5349 | 0.0403 (*) | 0.1659 |
| IL-9 | 229.7 ± 6.34 | 185.2 ± 5.28 | 191.8 ± 6.98 | 0.0003 (***) | 0.0007 (***) | 0.4525 |
| IL-10 | 12.72 ± 0.60 | 9.56 ± 2.14 | 8.57 ± 0.80 | 0.0669 | 0.0229 (*) | 0.6663 |
| IL-11 | 0.89 ± 0.90 | 2.76 ± 0.76 | 0.60 ± 0.38 | 0.0366 (*) | 0.6412 | 0.0301 (*) |
| IL-12/p40 | 4.94 ± 1.17 | 3.77 ± 1.87 | 4.04 ± 1.11 | 0.6013 | 0.7295 | 0.9722 |
| IL-12/p70 | 3.34 ± 0.59 | 3.54 ± 1.84 | 3.16 ± 0.98 | 0.9798 | 0.9831 | 0.9286 |
| IL-13 | - | - | - | - | - | - |
| IL-15 | 33.18 ± 2.24 | 27.81 ± 4.00 | 19.19 ± 2.23 | 0.1437 | 0.0027 (**) | 0.0268 (*) |
| IL-16 | 241.05 ± 61.39 | 382.7 ± 53.75 | 216.31 ± 32.86 | 0.0328 (*) | 0.8270 | 0.0165 (*) |
| IL-17 | 0.13 ± 0.10 | 0.14 ± 0.12 | 0.03 ± 0.01 | 0.9989 | 0.3746 | 0.3554 |
| IL-20 | - | - | - | - | - | - |
| IP-10 | 5.31 ± 0.69 | 48.57 ± 27.52 | 8.26 ± 2.97 | 0.0370 (*) | 0.9723 | 0.0488(*) |
| KC | 7.32 ± 2.72 | 13.73 ± 10.69 | 3.94 ± 1.01 | 0.4756 | 0.7970 | 0.2207 |
| LIF | 0.66 ± 0.18 | 1.58 ± 0.54 | 0.77 ± 0.09 | 0.0760 | >0.9999 | 0.2209 |
| LIX | 268.7 ± 27.41 | 229.2 ± 19.23 | 208.8 ± 36.03 | 0.6097 | 0.1840 | >0.9999 |
| MCP-1 | 3.81 ± 0.86 | 11.39 ± 9.57 | 1.49 ± 1.14 | 0.4260 | 0.9107 | 0.2105 |
| MCP-5 | 23.72 ± 7.91 | 224.4 ± 148.2 | 48.53 ± 16.07 | 0.0655 | 0.9345 | 0.1022 |
| M-CSF | 0.84 ± 0.20 | 0.85 ± 0.69 | 0.52 ± 0.10 | 0.9996 | 0.7363 | 0.7230 |
| CCL22 | 0.99 ± 0.23 | 5.03 ± 0.49 | 2.86 ± 0.44 | <0.0001 (****) | 0.0031 (**) | 0.0014 (**) |
| MIG | 36.13 ± 15.11 | 155.8 ± 100.7 | 52.99 ± 25.51 | 0.1134 | 0.9387 | 0.1751 |
| MIP-1a | 52.55 ± 9.31 | 36.41 ± 7.62 | 34.19 ± 7.56 | 0.1219 | 0.0800 | 0.9421 |
| MIP-1β | - | - | - | - | - | - |
| MIP-2 | 273.1 ± 31.23 | 244.4 ± 10.57 | 214.0 ± 11.89 | 0.2675 | 0.0272 (*) | 0.2366 |
| MIP-3α | 1.47 ± 0.06 | 1.05 ± 0.19 | 0.95 ± 0.02 | 0.0089 (**) | 0.0030 (**) | 0.5380 |
| MIP-3β | 31.22 ± 12.74 | 134.18 ± 16.86 | 79.63 ± 16.55 | 0.0005 (***) | 0.0204 (*) | 0.0119 (*) |
| RANTES | 2.41 ± 0.48 | 1.73 ± 0.12 | 1.34 ± 0.66 | 0.2614 | 0.0743 | 0.6069 |
| TARC | 0.72 ± 0.39 | 4.88 ± 1.90 | 2.60 ± 1.01 | 0.0163 (*) | 0.2433 | 0.1474 |
| TIMP-1 | 21.12 ± 18.29 | 1297.59 ± 443.64 | 476.21 ± 234.11 | 0.004 (**) | 0.2125 | 0.0307 (*) |
| TNFα | - | - | - | - | - | - |
| VEGF | 6.04 ± 1.69 | 12.49 ± 3.63 | 9.09 ± 2.62 | 0.0648 | 0.4203 | 0.3528 |
| 6Ckin/Exodus | 850.3 ± 210.8 | 955.1 ± 64.56 | 642.0 ± 101.7 | >0.9999 | 0.5391 | 0.2209 |
| TGF-β1 | 62.12 ± 60.71 | 87.47 ± 10.29 | 16.47 ± 4.25 | 0.6765 | 0.3278 | 0.0004 (***) |
| TGF-β2 | 8.89 ± 3.76 | 65.62 ± 28.09 | 26.01 ± 14.47 | 0.0214 (*) | 0.5262 | 0.0854 |
| TGF-β3 | 1.88 ± 0.43 | 19.13 ± 5.97 | 6.37 ± 3.56 | 0.0046 (**) | 0.4130 | 0.0190 (*) |
