## Supplementary material for "Clearance of senescent cells following cardiac ischemia-reperfusion injury improves recovery": r script file

library(plyr)
#load the input file containing z-scored (in rows, so individualy for each protein) peak area values
input<-read.delim("input.txt",stringsAsFactors = FALSE)
#specify the reference profile. The length should be equal to the number of samples
ref<-c(-1,-1,-1,1,1,1,-1,-1,-1)
N<-length(ref)
#generating ID for each row
input$ID<-1:nrow(input)


distance<-function(x,...){ #Calculates the distance between protein profile and reffference 
#(sum of suqred differences at each point/sample)
  rslts <- as.data.frame(matrix(NA,nrow=nrow(x),ncol=2))
  names(rslts) <- c("ID","r2")
  rslts[,"ID"]<- rownames(x)
  for(i in 1:nrow(x)){
    profile<-as.numeric(x[i,C])
    comp<-as.data.frame(cbind(ref,profile))
    comp$squares<-(comp$ref-comp$profile)^2
    rslts$r2[i]<-sum(comp$squares)
  }
  return(rslts)
}


p.threshold<-function(FDR,...){ # Estimates a maximum "distance" for a given FDR based on previously
#calculated permutation list
    p.th<-data.frame(matrix(unlist(perml)))
    p.th<-do.call(rbind, perml)
    p.th<-p.th[order(p.th$r2,decreasing = FALSE),]
    p.th<-p.th[1:(nrow(p.th)*FDR),]
    p.th<-p.th$r2[nrow(p.th)]
    assign(paste("FDR_",FDR,sep=""),p.th)

    return(p.th)
  }


pFDR.sign<-function(x, ...){ #Creates a new set of columns in the input file, 
  #indicating profile simmilarity at given FDRs (in this case 0.01 and 0.05)
  cnames<-colnames(x)
  C<-1:N
  y<-distance(x)
  x<-join(x,y)
  n<-ncol(x)
  p1<-p.threshold(FDR=0.01)
  p5<-p.threshold(FDR=0.05)
  for (i in 1:nrow(x)){
    if (x[i,n]<=p5){
      x[i, n+1]<-TRUE
    }
    if (x[i,n]>p5){
      x[i, n+1]<-FALSE
    }
    if (x[i,n]<=p1){
      x[i, n+2]<-TRUE
    }
    if (x[i,n]>p1){
      x[i, n+2]<-FALSE
    }        
  }
  colnames(x)<-c(cnames, "r2","pFDR.5","pFDR.1")
  
  return(x)
}

nperm=500 #Set number of permutations


#Here we generate a table of a set number of column permutations
for(i in 1:nperm){
  if(i==1){
    perm<-sample(1:N,N)
  }
  else{perm<-rbind(perm,(sample(1:N,N)))}
}
perml<-vector('list',nperm)


#Here we calculate the distances between protein profiles and the reference
for(p_ in 1:nperm){
  C<-as.numeric(perm[p_,1:N])
  perml[[p_]] <- distance(input, C)
} 

#Assesment of profile simmilarity
input<-pFDR.sign(input)

#Saving the result
write.table(input,"output.txt")
